# Deep Learning Assisted Mechanotyping of Individual Cells Through Repeated Deformations and Relaxations in Undulating Channels

**DOI:** 10.1101/2021.05.17.444390

**Authors:** Cody T. Combs, Daniel D. Seith, Matthew J. Bovyn, Steven P. Gross, Xiaohui Xie, Zuzanna S. Siwy

## Abstract

Mechanical properties of cells are important features that are tightly regulated, and are dictated by various pathologies. Deformability cytometry allows for the characterization of mechanical properties of hundreds of cells per second, opening the way to differentiating cells via mechanotyping. A remaining challenge for detecting and classifying rare sub-populations is the creation of a combined experimental and analysis protocol that would assure classification accuracy approaching 100%. In order to maximize the accuracy, we designed a microfluidic channel that subjects each cell to repeated deformations and relaxations. We also track the shape dynamics of individual cells with high time resolution, and apply sequence-based deep learning models for feature extraction. HL60 cells with and without treatment with cytochalasin D (cytoD), a reagent previously shown to perturb the actin network, were used as a model system to understand the classification potential of our approach. Multiple recurrent and convolutional neural network architectures were trained using time sequences of cell shapes, and shown to achieve high classification accuracy based on cytoskeletal properties alone. The best model classified the two sub-populations of HL60 cells with an accuracy of 95%. This work establishes the application of sequence-based deep learning models to dynamic deformability cytometry.

## Introduction

Cells’ characterization and classification are at the heart of understanding biology. In order to make the process biophysically and medically viable, both throughput and classification accuracy must be maximized. Classification accuracy on a single-cell level is especially important in identifying rare populations of cells such as circulating tumor cells^1^ or even cancer stem cells.^2^ These rare sub-populations can constitute a very small percentage (below 1%) of the whole population. The characterization based on physical properties of cells present an attractive, label-free basis for cells’ classification. These physical properties include optical properties like refractive index,^3^ electrical properties such as membrane capacitance,^4^ and mechanical properties like deformability i.e. the ability of a cell to change shape when an external force is applied.^5^ In particular, changes in mechanical properties have been linked to cellular differentiation,^6^ malignant transformation,^7^ formation of biofilms^8^ and even COVID-19 pathology.^9^ In this manuscript, we show the potential to classify cells with >90% accuracy based on mechanical properties alone.

Conventional methods for measuring the mechanical properties of cells include atomic force microscopy,^10,11^ optical tweezers^12^ and micropipette aspiration.^13^ While these methods are able to accurately measure mechanical properties of single cells, their throughput of roughly 1-10 cell(s) per minute is significantly slower than the ~ 10,000 cells per second achieved by most flow cytometers, which is needed to determine heterogeneities in cell populations and single cell classification.

In recent years, many microfluidic based methods have been proposed for measuring the deformability of cells with significantly improved throughput, closing the gap between flow cytometry and mechanical phenotyping methods. Microfluidic approaches to characterize mechanical properties of cells offer a label-free and high-throughput characterization that is cost-effective and scalable. In one class of approaches, cells are squeezed through constrictions smaller than a cell’s diameter.^14–17^ Here, the deformability is often based on the transit time through the channel, measured by recording electrical impedance. These methods typically lack optical characterization; thus, they are unable to develop a correlation between the measured electrical parameters and actual changes to the cell shape that occur due to deformation. In addition, using narrow channels that induce strong deformations of the cells compromises cells’ viability.

In another group of microfluidic methods, cells are deformed by the hydrodynamic forces formed inside the channel that is larger than the cells, as done in extensional deformability cytometry (xDC).^18^ xDC utilizes a cross channel design to deform and image cells at flow rates reaching ~1000 μL min^−1^. This approach has been used to classify malignant pleural effusions, differentiate multiple stem cells, as well as identify transitions in the cell cycle. Real-Time Deformability Cytometry^19^ (RT-DC), based on a straight, narrow microfluidic channel, has emerged as another promising method for high-throughput and label-free phenotyping of individual cells. RT-DC uses lower flow rates than xDC (~1 μL min^−1^), and was demonstrated to be sensitive to perturbation of cellular actin networks. RT-DC quantifies deformability based on a steady-state image of each deformed cell captured at the end of the microfluidic constriction. While the image of the steady state shape is useful for extracting a cell’s mechanical properties such as Young’s modulus, the single image misses the shape dynamics that accompany the deformation. Because these dynamics are rich in mechanical information, recording them has the potential to expand the available markers for mechanical phenotyping.

Deformation dynamics together with morphology based features were used in expanding the available feature space for classification in xDC.^20^ With the use of support vector machines (SVM), the addition of morphology based features indeed led to improved classification; however, these features could not be used to phenotype populations which differed only in cytoskeletal properties. Further, the measured deformation dynamics from this approach had only a small contribution to the gain in classification accuracy, indicating that time-dependent viscoelastic properties were not being fully probed in these measurements. Another method, dynamic RT-DC (dRT-DC),^21^ proposed recording the time series of deformations a single cell undergoes until it reaches a steady state shape, and enabled determination of multiple viscoelastic properties. These measured quantities showed some improvement in classification when using a logistic regression model, however, the step-stress induced by the straight channel precluded the ability to probe frequency dependent components of the cytoskeleton, leaving unused mechanical information.

The classification algorithms, such as SVM and logistic regression, used by all of the above mentioned methods are heavily dependent on human derived features, such as size, deformation, and brightness. To automatically learn the mechanical and morphological features, a dense neural network (DNN) has recently been applied to deformability cytometry images, which classifies cells using the images directly.^22^ There remains a large potential to apply deep learning models, in particular sequence based neural networks, to mechanotyping of cells using shapes alone. Doing so provides a path to expand upon the currently used mechanical markers, which would be useful in classifying cell populations in which mechanics, but not morphology, are perturbed.

In this manuscript we maximize the classification between cell populations differing only in cytoskeletal properties through the introduction of a multiple deformation microfluidic channel, tracking of shape dynamics, and application of sequence based deep learning models for feature extraction. We utilize a microfluidic channel whose shape was designed to subject each cell to multiple deformations and relaxations through hydrodynamic forces. The channel has an undulating width and contains a cavity flanked by two narrower regions.^23^ Two sub-populations of HL60 cells before and after treatment with cytoD were used as a model system to probe classification potential of our method. CytoD was found to perturb actin filaments with minimum effect on the cells’ shape or morphology when in suspension. As a cell enters the first narrow zone, it undergoes the first deformation becoming elliptically shaped. Upon entering the cavity, the cell relaxes to its original spherical shape, becoming elliptical again in the second narrow zone. As the videos are recorded with the time resolution of at least 11,000 frames per second, we can observe the deformation dynamics with high precision, revealing quantitative insights into the deformation/relaxation times. We show that using these multiple features as well as time series of deformations, one can significantly increase the accuracy of classification of HL60 cells before and after cytoD treatment. Specifically, we achieve an accuracy of 70% when using an individual feature such as maximum deformation, while taking advantage of the shape dynamics increases the classification accuracy to >90%, which is greater than the accuracy obtained from studies using both biomechanical and bioelectrical features for the same cell model system.^24^

The increased accuracy of cells’ classification was made possible by subjecting the time-based sequences of cells’ shapes to deep learning models such as recurrent neural networks (RNN). The deep learning methods utilizing time-based sequences of features showed an increase in classification accuracy compared to traditional machine learning methods, and were crucial in achieving >90% accuracy. Most importantly, a convolutional neural network (CNN) was used in conjunction with an RNN to utilize sequences of binary masks as input features. We also applied Shapley values known from game theory to probe which deformability parameters our experiments provide contributed most to the classification accuracy.

We envision this channel design, along with the demonstrated deep learning models to enhance cell classification and ultimately sorting. While the deep learning characterization and classification are currently performed post-measurements, similar deep learning models with proper hardware can currently process ~2,000 fps, leading to the possibility for in situ sorting.

## Results and Discussion

### Channel Design and Data Acquisition

We designed a microfluidic channel that subjects individual cells to repeated compressions and relaxations. The multiple deformations of the cells are created by the channel shape, specifically the presence of a cavity and two narrow zones. The varying channel width is expected to create inhomogenious velocity profiles leading to temporal changes in the cells’ shape.^25^ To test this channel design for mechanotyping, we pumped a suspension of HL60 cells (ATCC-240) in methylcellulose (1% w/v) solution through a channel of consecutive constrictions of width 25 μm, 50 μm, and 25 μm at a flow rate of 1 μL min^−1^. Sheath flow focusing was used to ensure that all cells passed through the channel along its center axis and experienced the same forces. The microfluidic chip was placed on a 10x magnification inverted microscope. To enable probing dynamics of cells’ shape at this high flow rate, we customized the microscope to allow for high speed imaging. We replaced the light source with a high-powered LED array and accompanying 3D printed mount (Figure 1a). A high-speed camera was adapted to fit the microscope and videos of translocating cells were recorded at 11,000 frames per second, revealing the shape dynamics as a cell moves along the channel axis.

**Figure 1:**
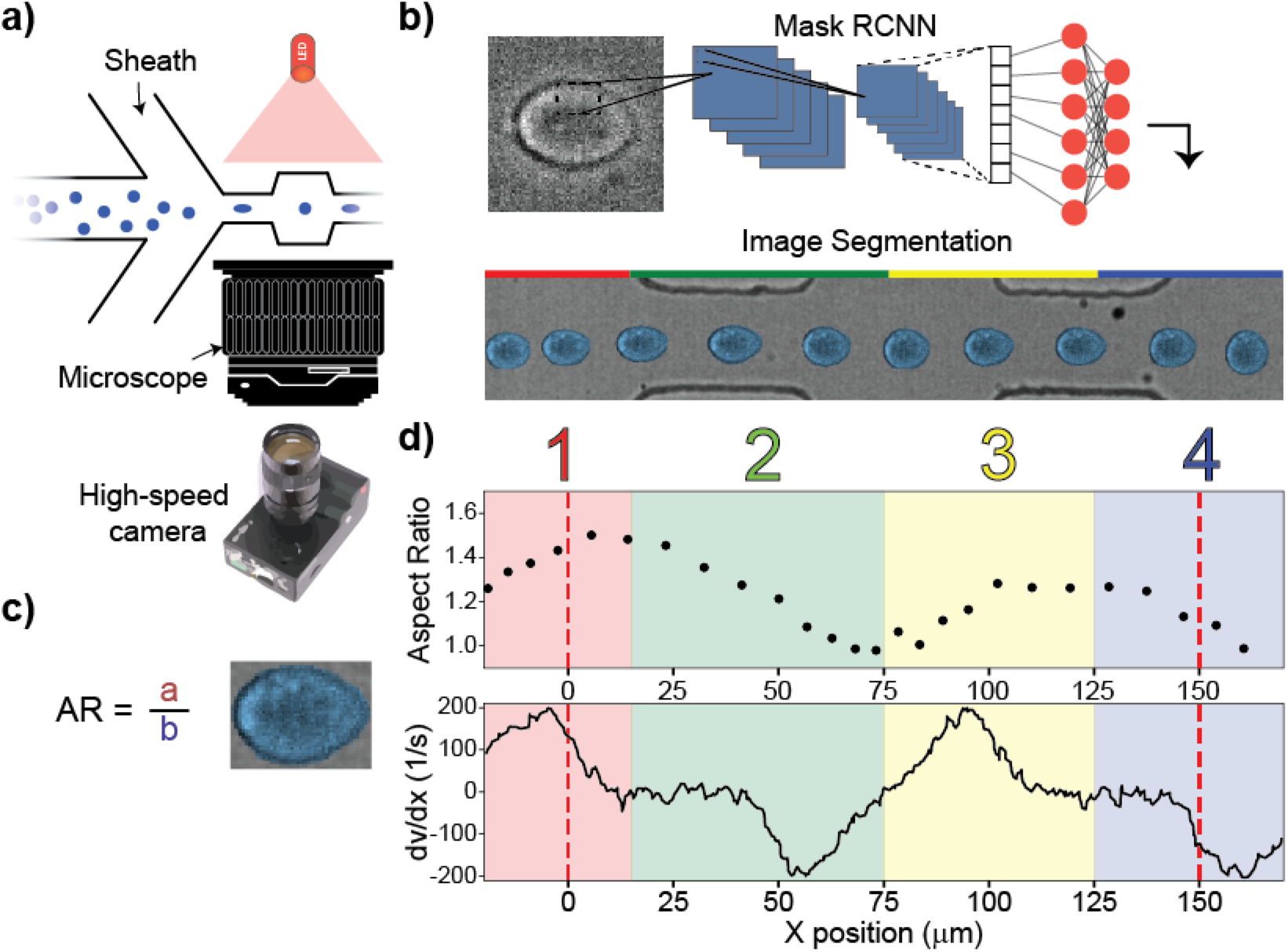
Principles of repeated mechanotyping. a) Channel design utilizing sheath flow, a high-powered LED and a microscope. The microfluidic channel that enables characterization and classification contains a cavity placed between two narrow zones. b) Data is captured by a high-speed camera, creating videos at 11k fps. Cell borders are detected and fit using Mask-RCNN. c) The cell deformation, AR, was quantified as the ratio of two axes of an ellipse that approximates the cell’s shape. (d) (Top) The aspect ratio versus position relative to channel entrance of a single cell as it passes through the channel. (Bottom) COMSOL simulation showing the derivative of velocity vs channel position, which is proportional to the shear stress.

After videos had been collected, the data was processed with a number of python scripts (Supplementary Figure 2). The first script encompasses a lightweight CNN that filters out empty frames, and reduces the amount of data to be processed by ~80%. Next we employed a modified architecture of the Matterport^26^ implementation of Mask-RCNN,^27^ which is used to detect instances when objects are in an image and the masks associated with each object. Our version of Mask-RCNN is trained on 500 hand-labeled images of cells with varying levels of focus (Supplementary Note 1). The masks detected by this network are then processed using a custom tracking algorithm to find trajectories of individual cells (Supplementary Note 2). Figure 1b shows subsequent snapshots of one cell as it passes through the channel. We found that the undulating channel design results in complex dynamics of the cell’s shape, and leads to regions of differing deformations. Specifically, the cell underwent a strong deformation at the entrance of the channel and in the first narrow constriction; the cell then relaxed to a spherical shape in the cavity, and started to deform again when approaching the second narrow constriction.

We defined the deformation as the aspect ratio, AR, of the two axes of the ellipse: the axis parallel to the channel axis and the perpendicular axis (Figure 1c). AR = 1 corresponds to a sphere, whereas AR>1 corresponds to an extension along the channel axis. Figure 1d summarizes how the magnitude of AR evolves as the cell shown in Figure 1b passes through the channel. In order to qualitatively understand the deformation trace, we ran a computational fluid dynamics simulation with COMSOL multiphysics in a cell-free undulating channel using the Navier-Stokes equations with creep flow at experimental flow rates. Shear stress in different parts of the channel can be analyzed through the derivative of velocity in the center of the channel with respect to axial position shown in Figure 1d. The cells experience a large velocity gradient at the entrance of the channel, leading to large stresses and deformations. For the single cell trace shown in Figure 1d, we observe a peak deformation of AR = 1.49 in the first region (R1), marked in red. The velocity gradient then reaches a steady state value within the first narrow constriction, (10 μm - 50 μm) and the cell deformation decreases. Before the cell can reach a steady state deformed shape, it enters the cavity where the velocity gradient begins to decrease and then again rapidly increases. The cell returns to a spherical shape, AR = 1, where dv/dx = 0, which occurs at ~75 μm. The cell then begins to deform again due to the velocity gradient at the entrance of the second narrow constriction, reaching a second peak deformation of AR = 1.30 in region three (R3). The maximum deformation observed in the second constriction is lower than the maximum deformation measured in the first constriction. We hypothesize it is because the cell is subjected to shear stress over a relatively short time and distance when transitioning from the cavity to the second narrow region, as compared to the transition from the bulk channel to first narrow constriction entrance. The distance from the middle cavity to the position of peak deformation in the second narrow region is only 25 μm, whereas at the initial inlet the cell begins deforming from shear stresses ~50 μm away from the entrance. Figure 2a shows values of AR for ~700 HL60 cells examined in the same conditions. The same trend for all cells has been observed: the cells reached the maximum deformations in the first narrow zone, relaxed to a sphere in the cavity, and underwent another deformation in the second constriction.

**Figure 2:**
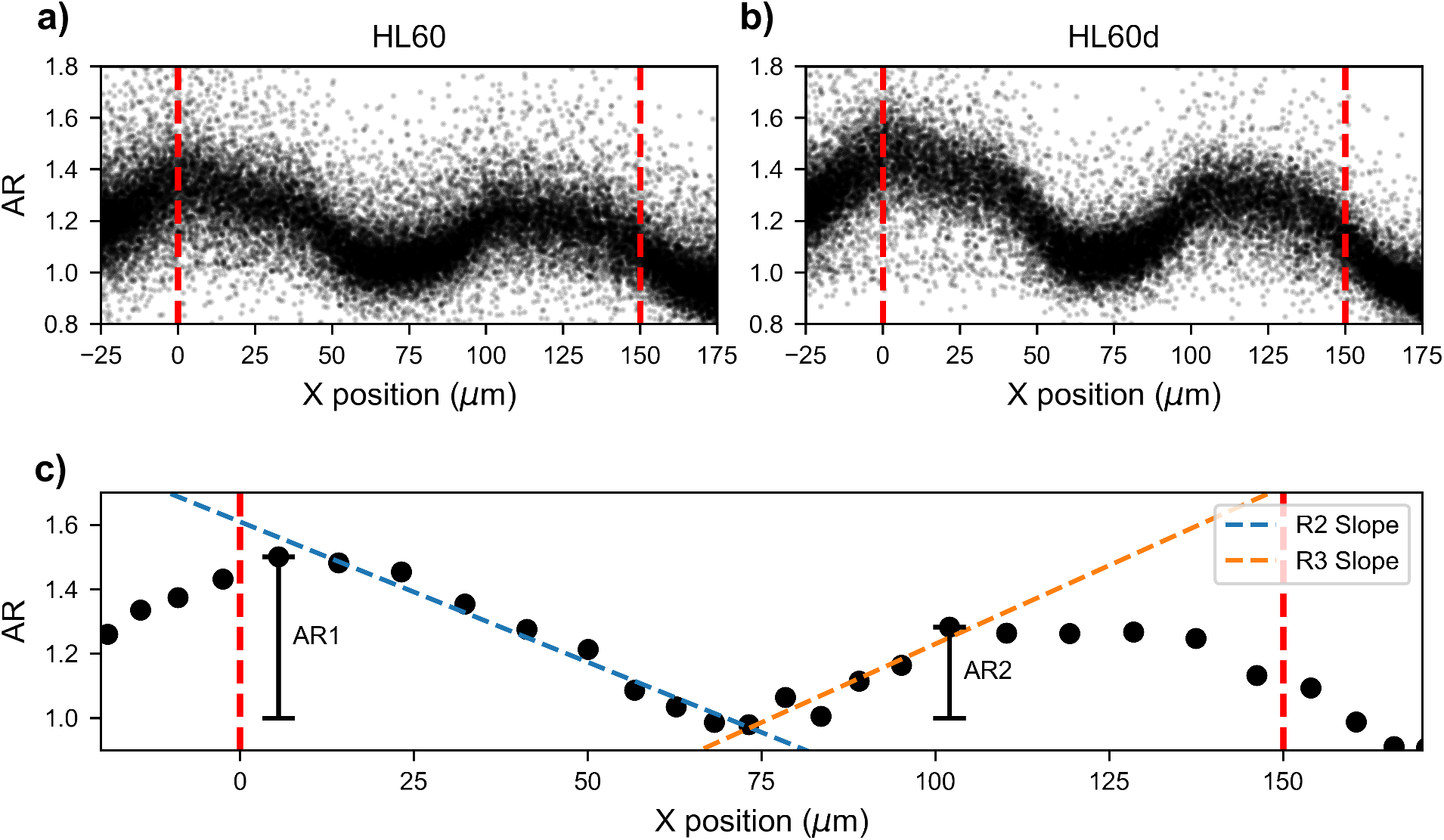
Single cell deformation traces. Deformation dynamics are shown for single cells translocating through the channel. Deformation is defined as the aspect ratio of two axes of an ellipse, and the position along the channel axis is determined by the centroid of the mask. The CytoD treated HL60 cells show increased deformation versus untreated HL60 cells in narrow regions of the channel. Both cells experience a smaller maximum deformation in the second narrow region, as compared to first narrow zone. Channel inlets and outlets are marked by red dotted line. Deformation and relaxation occur twice within the channel. a) Full population of single cell traces of aspect ratio versus position for untreated HL60 cells. b) Full population of single cell traces of aspect ratio versus position for cytoD treated HL60 cells. c) Single cell example of parameters that are determined: maximum AR values in the two narrow zones as well as relaxation and deformation slopes.

Taking advantage of the time-series of cells’ positions and shapes, we also quantified the dynamics of deformation. To this end, we used a linear model to fit the trace from the peak deformation in the first narrow zone to the position where the cell relaxes to a spherical shape, resulting in a slope of −8.71 × 10^−3^ μm^−1^. A similar analysis can be performed for the cell entering the second narrow zone by fitting a line between the spherical shape in the cavity to the maximum deformation in the second narrow zone, obtaining a value of 9.75 × 10^−3^ μm^−1^. An example of these parameter fits to a single cell trace can be seen in Figure 2c. Recordings shown in Figure 2a allow determination of the slopes for all examined individual cells.

### Application of the undulating channel to probe perturbation of actin networks

A recent study comparing different microfluidic methods for measuring deformability suggested there is a threshold of shear rate beyond which changes to the actin filaments in the cytoskeleton are not detectable.^28^ To test whether our method, with a flow rate of 1 μL min^−1^, is sensitive to the cytoskeleton changes, we created a sub-population of the HL60 cells with perturbed actin networks. We used cytochalasin D (cytoD) to disrupt actin polymerization,^29^ previously shown to increase deformability. CytoD treated and untreated HL60 cells have been previously used as a model system to evaluate mechanotyping techniques.^19^ With the cells suspended in methylcellulose, we separately passed each population of cells through our microfluidic channel.

Figure 2b shows subsequent deformations of HL60 cells after treatment with cytoD. A visual inspection of the aspect ratios in the first and second narrow constrictions suggests that the cytoD treated cells deform more than the untreated cells. This finding is in agreement with earlier reports that used the same cell lines and also observed increased deformability of the cytoD treated sub-population.^30^ Since the time series of shapes is recorded with a high-speed camera and consist of ~30 frames for each cell, we can perform a comparative statistical analysis not only of the deformation, AR, but also the slopes that quantify the deformation dynamics, R2 and R3 in Figure 2c. The R2 slope is a measure of relaxation dynamics, while the R3 slope describes deformation. Figure 3 shows the AR in the two narrow zones and the two slopes for the two sub-populations of the HL60 cells: untreated and cytoD treated cells. The magnitudes of AR are consistently higher for the cytoD treated sub-population, which confirms that the method is sensitive to actin network perturbations. Interestingly, the slopes that represent deformation dynamics are greater for cytoD-treated cells, suggesting that these cells are more responsive to external forces. It is important to note that the mean diameter of the two populations is nearly identical (Supplementary Figure 3), confirming the cells experienced the same forces.

**Figure 3:**
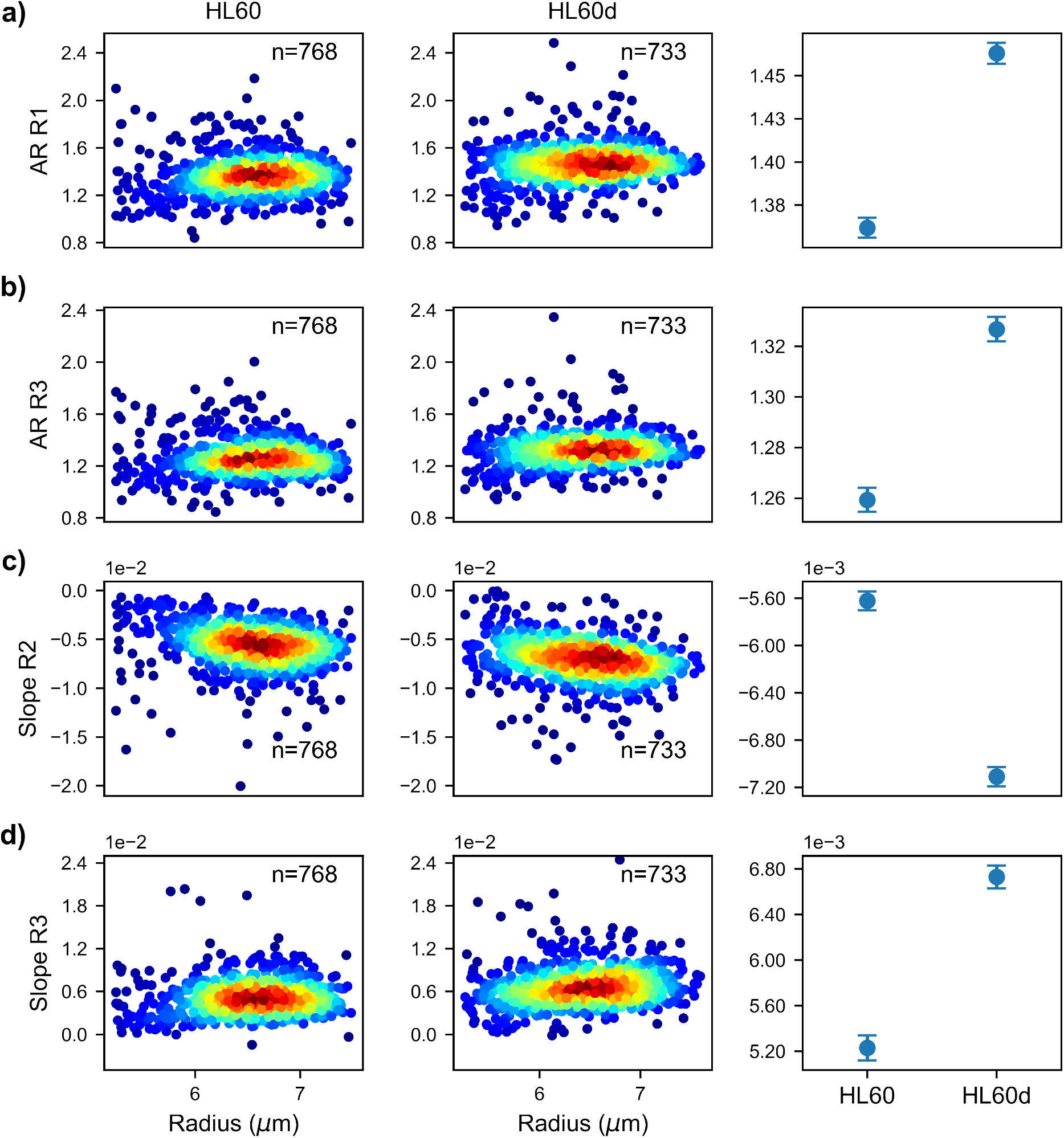
Comparison of measured deformability features between untreated and cytoD treated HL60 cells a) Heat map of maximum aspect ratio in first narrow zone (R1) for both untreated and cytoD treated cells. Mean of each population is reported where the reported error is standard error of the mean for all subsequent calculations of the mean. b) Heat map of maximum aspect ratio in second narrow region (R3) c) Linear fit slope from maximum deformation in first narrow region (R1) to relaxation to sphere in cavity (R2). d) Linear fit slope from relaxed state in cavity (R2) until maximum deformation in second narrow region (R3).

Though we can measure differences in the means of the two populations’ deformation, no single feature alone can provide accuracy > 70% for single cell classification (Supplementary Note 4). In order to be able to discriminate between the two cell types on a single cell basis, multiple features describing the cell deformation (Figure 3) are required to be considered simultaneously.

### Feature Extraction and Machine Learning Model Comparison

To improve classification accuracy and to better understand the cells’ deformations, we extracted features (Supplementary Note 4) to be used in classification models that in turn would enable probing classification potential of our microchip design. We first investigated the performance of traditional machine learning models to distinguish the two cells’ sub-populations, HL60 untreated and cytoD treated cells. The models we chose have been previously employed for classifying cytometry data and include: k-nearest neighbors^31,32^ (KNN), SVMs,^20^ random forests^33–35^ (RF) and logistic regression^36,37^ (LR). All models were implemented using scikit-learn and all relevant hyperparameters were optimized (Supplementary Figure 5).

All models showed relatively good performance in recognizing untreated and cytoD treated cells (Figure 4a). To determine the features’ impact on the RF model output, we utilized Shapley values that helped us understand how the channel design generate informative features that enable cells’ classification. Shapley values were developed for coalitional game theory and inform one how to fairly attribute success to the constituent parts. In order to make use of this method, the SHAP python library^38^ was employed in conjunction with the trained RF model to attribute overall model performance to individual features (Figure 4b). Shapley values show that the linear fit to the relaxation process occurring in R2 and maximum value of AR in R1 have the most weight in deciding classification between the two cell classes. The introduction of Shapley values provides interpretability and demonstrates that the region from peak AR to the relaxed state contains the most information for classification. The analysis also revealed the importance of another temporal feature, the R3 slope, in the classification (Figure 2c). Based on these observations, we hypothesized that classification could be further improved by incorporating the sequential nature into the model design. We then decided to explore deep learning approaches to create a model that can extract shape dynamics.

**Figure 4:**
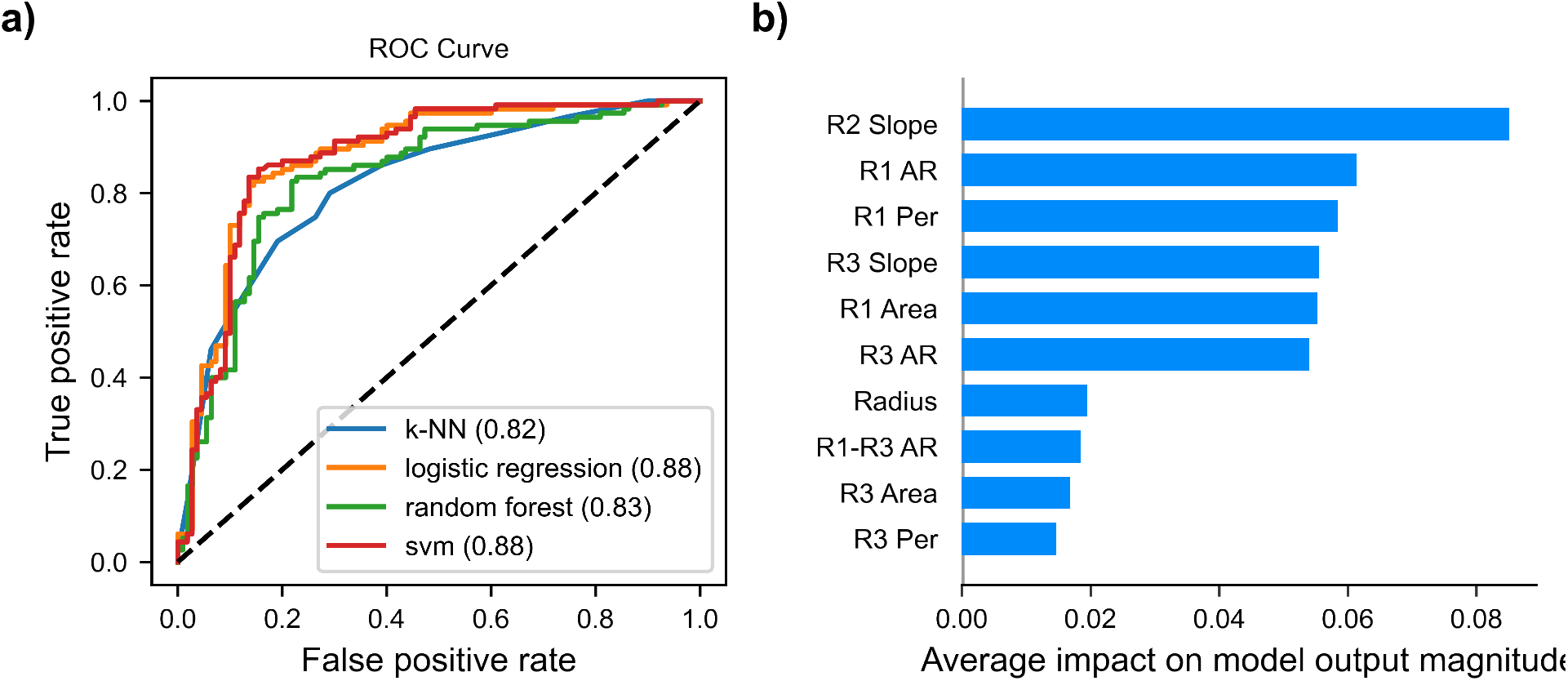
a) Receiver operator characteristic (ROC) curve for traditional machine learning models. Ideal curves approach top left and exhibit maximized area under the curve (AUC). Performance obtained by guessing shown as black dotted line. AUC values are shown in parentheses. b) SHAP feature importance plot obtained using trained RF model.

### Deep learning for enhanced classification

RNNs are considered the optimal tools for handling sequential data^39^ and are often used for translation and sequential prediction tasks. In short, RNNs function by receiving a single input from the full sequence, process it and feed the output into a copy of itself along with the next time step. We implemented a variant of RNNs, gated recurrent units (GRUs) and extracted sequential features, such as aspect ratio, from the data that could be used as inputs in these models (Supplementary Note 2). Though other deep learning models (Supplementary Note 6) were tested using these data, GRUs showed the best classification performance. While the observed performance is an improvement (Figure 5) compared to the SVM, there are limitations in using hand-selected features that could potentially hinder the performance. We therefore sought to improve accuracy by using the binary masks output by the segmentation network as inputs. We decided to employ CNNs, which were originally developed for video recognition^40^ and thus are ideally suited for extracting features from images. In order to enable extracting temporal patterns from the images, GRU layers were added on top of the CNN layers. A schematic diagram of the model is shown in Figure 5a. The inputs for the model are the masks output by the segmentation network in addition to ellipses fit to the mask. In order to ensure that the HL60 cells before and after cytoD treatment intrinsically do not exhibit morphological differences, a control study was done. To this end, a CNN was trained to separate the two HL60 sub-populations using masks from the channel cavity. The best validation accuracy attainable was 65%, indicating poor classification potential of the morphology (Supplementary Note 7).

**Figure 5:**
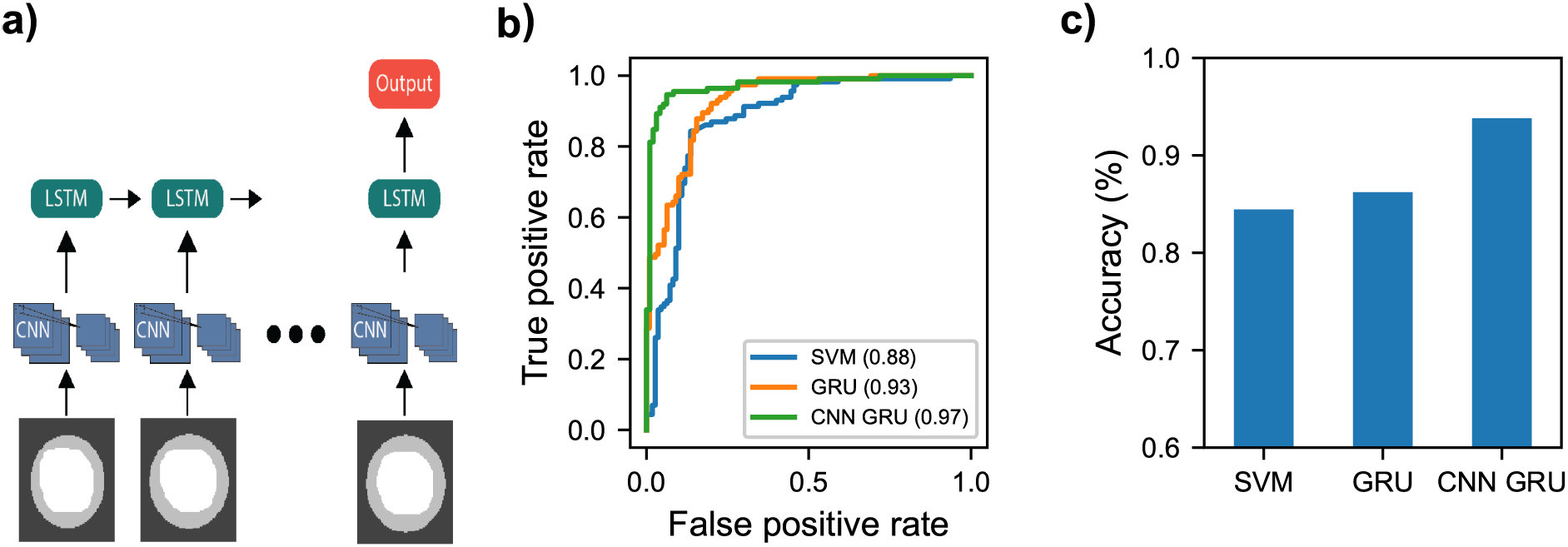
a) General flow of CNN-GRU. Sequences of masks are padded and used as inputs. CNN and GRU layers use identical weights for each time step. b) ROC curve for SVM, GRU and CNN-GRU using the validation set. AUC shown in parentheses. c) Plot showing model accuracy on validation set.

While CNN-RNNs traditionally use raw frames as inputs, here our input complexity is drastically reduced since segmentation was already performed. Our CNN-GRU architecture was refined by searching different filter sizes for the CNN layers, changing the number of dense and GRU layers, as well as adjusting the dropout rates. The optimized model (Supplementary Note 6) shows superior performance to all other models tested (Figure 5b, c). Our particular model shows the highest ROC AUC (0.97) of all models tested. To investigate the potential of over-fitting, 5-fold cross-validation was performed in addition to the use of a validation set. Similar results were obtained for each training regime (Supplementary Note 8). The high validation set performance shows the richness of the deformability dynamics and their potential to aid in classification.

## Conclusions

In summary, we show that the application of a channel with an undulating width reveals a multitude of information on cells’ mechanics that otherwise is not accessible. The channel we designed induces multiple velocity gradients along its length, and consequently the cells undergo multiple deformations and relaxations. We tracked the transient shape of cells as they translocated through the channel with Mask-RCNN and measured the deformation multiple times at each channel region. These measurements allow one to find peak deformation as well as deformation and relaxation dynamics that were used then as inputs into linear-boundary machine learning models to discriminate between HL60 and actin polymerization inhibited HL60 cells. One of these models enabled the use of Shapley values which highlighted time-based features, directing future model construction. Neural networks such as GRUs were then employed to learn features from the timeseries deformability data. The application of GRUs resulted in improved classification compared to the traditional techniques. Finally, we extended the GRU model by using sequences of binary masks as inputs, and utilizing a CNN layer to extract shape based features. The CNN-GRU model resulted in the highest classification accuracy of the two cell populations that reached 95%. The improved accuracy of cells’ classification is expected to become the basis for in-situ characterization and sorting, especially for important for segregating rare cells, including circulating tumor cells^1^ and even cancer stem cells.^2^

## Methods

### Cell Culture

HL60 (ATCC CCL-240) cells were cultured in suspension with Iscove’s Modified Dulbecco’s Medium (IMDM) supplemented with 10%(v/v) FBS(Fisher brand) and 1% (v/v) penicillin-streptomycin and maintained in an incubator at 37°C with 5% CO_2_. Cells were passaged through dilution every 2-3 days to maintain a density between 10^5^ and 10^6^ cells mL^−1^. Prior to experiments, cells were centrifuged at 112 relative centrifugal force (RCF) for 5 minutes and resuspended to concentrations of 2-3×10^6^ cells mL^−1^ in PBS with 1% (w/v) methylcellulose (Spectrum 4,000CP).

To create a populations of HL60 cells with perturbed actin networks, cells were incubated for 10 minutes in 1 μM cytochalasin D (cytoD) (Sigma) which had been diluted 10x from stock solution with dimethyl sulfoxide (DMSO).^30^ Cells were spun down at 112 RCF for 5 minutes and resuspended in 1% (w/v) methylcellulose solution.

### Channel Preparation and Data Acquisition

Microfluidic channels were prepared with a master mold using standard photolithography with negative SU-8 photoresist. SU-8 photoresist (Kayaku Advanced Materials Inc.) was dispersed on 50 mm wafer using laurel spinner to a thickness of 20 μm. The wafer was subsequently exposed and baked to form the master. 184-Sylgard polydimethyl siloxane (PDMS) was then pipetted over the SU-8 master and baked for ~4 hours at 75°C. Channel inlets and outlets were created with a 1.5 mm biopsy punch. PDMS channels were cleaned and dried with isopropyl alcohol, methanol and water before being bonded to a glass cover slip using a corona discharge wand (ETP). Bonded PDMS/glass samples were heated for an additional hour at 90 °C to promote further adhesion.

Prepared cells suspended in methylcellulose solution were pumped through a microfluidic channel using a Genie-plus syringe pump(Kent Scientific) at a rate of 1 μL min^−1^. Cells were focused laterally in the channel using a sheath flow geometry with a flow rate of 2 μL min^−1^. The sheath and core flow were allowed to equilibrate for 10 minutes before data was taken. Cells were illuminated using a high powered LED and imaged at 10x magnification in brightfield on an inverted microscope. A Chronos 1.4 high-speed camera (Krontech) imaged passing cells at a frame rate of ~11,000 fps with 1 μs exposure time and a resolution of 880×140 to encapsulate the full channel length. The size of each pixel is 0.26μL/pixel. Maximum blurring induced by cell movement is ~ 0.1 μL. 8 second videos were recorded and saved which require ~15 minutes to offload from camera memory. Experiments were conducted across two biological replicates and three technical replicates.

### Detection, segmentation, and tracking

A CNN was trained to identify frames with cells. Frames labeled as containing cells were then passed to the Mask RCNN.MASK-RCNN network was used to segment cells from images and fit masks. The Matterport implementation^26^ was used with Tensorflow 2.2. The network was trained using a NVIDIA 1070TI on a hand labeled dataset of ~300 images of HL60 and HL60d cells across multiple independent experiments using VGG Image annotator 1.0. The network was initialized with weights from the COCO dataset and the architecture was modified in order to increase the output resolution of the predicted masks. The training schedule consisted of training head layers for 20 epochs at a learning rate (LR) of 10^−3^, 50 epochs training 4+ layers at LR/10, and 50 epochs training all layers at LR/10. Training curves and further details can be found in the Supporting Information file 1.

### Calculation of features

Detected events with impossible trajectories (i.e. no event will begin in the middle of the channel) were discarded. To ensure the fits are accurate, a convex hull was fit to the cell mask and events were filtered out where a single frame had a ratio ¿ 1.1 between the original and convex hull fit. Detected particles with radii equal to three standard deviations from the mean were not included in analysis, as they often contained cell/channel debris, or clumps of multiple cells.

Shapes were acquired and assigned to each cell as a function of time and position in the channel. Time is determined by the frame rate of the camera and position was determined from finding the centroid of each mask. The channel was manually staged to place the (0,0) coordinate as the center entrance of each channel. Pixels were converted to microns using a ruler.

Ellipses were fit to the detected masks and the aspect ratio of the ellipse was used to describe the cell deformation, AR. Aspect ratio in the two narrow constrictions were determined by the maximum achieved aspect ratios in the respective regions. R2 slope was calculated by fitting a linear model from the position of R1 AR to the position where the cell assumes the most spherical shape in the cavity (~ 75 μm). R3 slope was calculated by fitting linear model from the circular postion to the position of maximum deformation in the second narrow constriction. A full description of parameters can be found in SI (4).

### Comsol Simulation

A finite element simulation was conducted using Comsol Multiphysics 5.3 to simulate the velocities and stresses experienced in the undulating channel. A simplified 3-D model of the channel was modeled in the laminar flow module using the Navier-Stokes equation with creep flow in steady-state. An extra fine meshing was chosen for the area near the narrow section of the channel and normal meshing was chosen for the reservoir.

### Traditional machine learning model training

All models were created and trained using python 3.6 and scikit-learn. The features used are listed below. The data were shuffled and split according to the following ratio: 70:15:15 for train, test and validation respectively. A function was built for the standardization based on the train data and then used to transform the validation and test data so as to not leak information. Receiver operator characteristic (ROC) area under the curve (AUC) and accuracy are reported using the test data.

### Sequential Scalar Data Preprocessing

Roughly 1500 sequences of aspect ratio, perimeter, deformability and area were selected as inputs. Data previously filtered were first aligned so that all sequences started and ended at a true x-position of −30 and 170 respectively. Data were then padded to length 50 and then shuffled and split into training, validation and test partitions with relative sizes 70%, 15%, 15% respectively.

### Image Data Processing for Sequential Models

For sequential models using image data, masks were first cropped to 90×90. An ellipse fitted to the mask shape was created using scikit-image and added to the second channel. Each sequence of these two-channel images were padded to length 50. The data were shuffled and split according to the following ratio: 70:15:15 for train, test and validation respectively.

### CNN-GRU

Sequences of images were used as inputs for a CNN-GRU with three CNN layers, two GRU layers and one output dense layer. The hyperparameters were optimized over 175 epochs using the test set. Model performance was assessed using a hold-out validation set, in addition to 5-fold cross-validation. Further details can be found in the Supporting Information.

### Code Availability

The python scripts implemented for data processing and training machine learning models are available at: https://github.com/siwylab/time-series-dc.

## 1 Data availability

The data that support the findings of this study are available from the corresponding author upon reasonable request.

## 2 Acknowledgment

The work was supported by UC Cancer Research Coordinating Committee, C21CR2129. This material is based upon work supported by the National Science Foundation under grant number 1633631. This work was funded by an opportunity award from the UCI Center for Complex Biological Systems, through NIH-NCI U54-CA217378.

## 3 Contributions

Z.S., C.C. and D.S conceived the study. C.C. and M.B. created experimental setup and protocols. C.C. and D.S. performed experiments on HL60 and Hl60d cells. D.S created deep learning models for classification. C.C. and D.S. performed data analysis. Z.S., C.C. and D.S wrote the manuscript. Z.S., M.B., S.G. and X.X. assisted in data interpretation. M.B., S.G. and X.X provided edits and revisions. X.X. provided guidance on the machine learning component. Z.S. supervised the project. All authors reviewed the manuscript.

