## Supplementary Information for "Deep Learning Assisted Mechanotyping of Individual Cells Through Repeated Deformations and Relaxations in Undulating Channels"

### Supplementary Note 1
### MASK-RCNN Training

MASK-RCNN network was trained to segment cells from images and determine masks. Network was trained on hand labeled dataset of HL60 and HL60d across multiple independent experiments using VGG Image annotator 1.0. MASK-RCNN is able to accurately fit masks at various focusing and lighting conditions with a mAP and mIOU $> 0.9$, reducing bias in overall mask fitting between experiments. Matterport implementation which was updated to run on Tensorflow 2.2 was used. The training schedule consisted of training head layers for 20 epochs at a learning rate (LR) of $10^{-3}$, 50 epochs training 4+ layers at LR/10, and 50 epochs training all layers at LR/10.

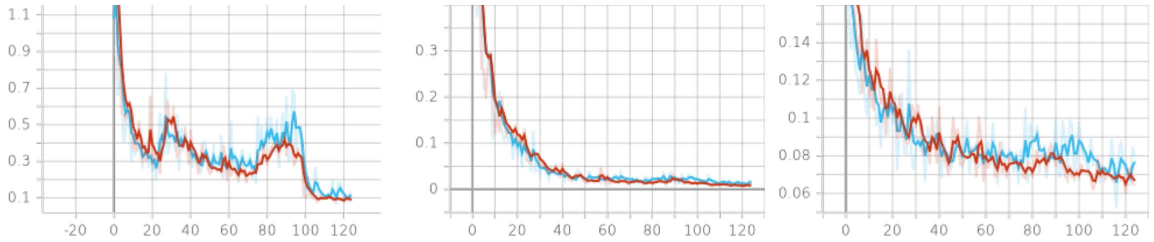


**Figure S1:** Training curves for Mask-RCNN. a) Total loss per epoch. b) Bounding box loss per epoch. c) Mask loss per epoch.

### Supplementary Note 2
### Python scripts for data processing

The python scripts implemented for data processing and training machine learning models are available at: https://github.com/siwylab/time-series-dc.

### Supplementary Note 3
### Comparison of radii between classes

We measure the radii of cells by measuring the area and perimeter of cell masks where shear stress is zero, which occurs at 75 μm. The radii is the average of fitting a circle with equivalent area and perimeter respectively. We find a small discrepancy between the mean radii of HL60 cells and HL60 cells treated with cytoD. Cells with radii larger than $3\sigma$ from the mean were removed from analysis, as they were often small cell or PDMS debris, or clumps of two or more cells.

### Supplementary Note 4

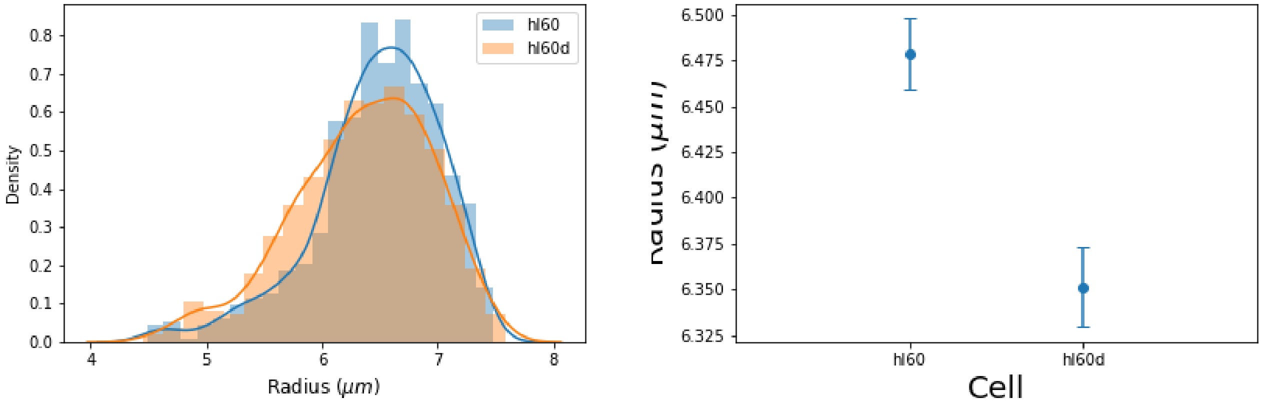


**Figure S2:** HL60 and HL60d radii comparison. a) Distribution of radii for two cell populations. b) Error bars represent standard error of the mean.

### Feature Table

| Feature | Description | Classification Accuracy |
|---|---|---|
| Radius | Radius of cell in cavity where deformation is minimal. Average radius of circle of equivalent perimeter and circle of equivalent area. | 57% |
| R1 AR | Maximum aspect ratio of fitted ellipse measured in first narrow constriction. | 71% |
| R3 AR | Maximum aspect ratio of fitted ellipse measured in second narrow constriction | 69% |
| R2 Slope | Slope of linear fit between position of maximum deformation in first narrow constriction and position of least deformation in cavity. Represents cell relaxation. | 71% |
| R3 Slope | Slope of linear fit between position of least deformation in cavity and position of maximum deformation in second narrow constriction. Represents cell compression. | 64% |
| R1-R3 AR | Difference between AR1 and AR2 | 59% |
| R1 Per | Average perimeter of mask in first narrow constriction. | 59% |
| R3 Per | Average perimeter of mask in second narrow constriction | 57% |
| R1 Area | Average area of mask in first narrow constriction | 60% |
| R3 Area | Average area of mask in second narrow constriction | 57% |

### Supplementary Note 5

#### Classical machine learning model training

All models were created and trained using python 3.6 and scikit-learn. The features used are listed below. The data were shuffled and split according to the following ratio: 70:15:15 for train, validation and test respectively. A function was built for the standardization based on the train data and then used to transform the validation and test data so as to not leak information. Receiver operator characteristic (ROC) area under the curve (AUC) and accuracy are reported using the validation data.

**Logistic Regression Classifier** - The l2 regularization was optimized through a grid search from 0.1 to 1.5 with 10 values tested.

**k-Nearest Neighbors** - The leaf size and number of nearest neighbors were refined in a grid search.

**Support Vector Machine** - The regularization strength, kernel, and degree for the polynomial kernel were optimized in a grid search.

**Random Forest** - Using the gini criterion, different depths were tested in a grid search.

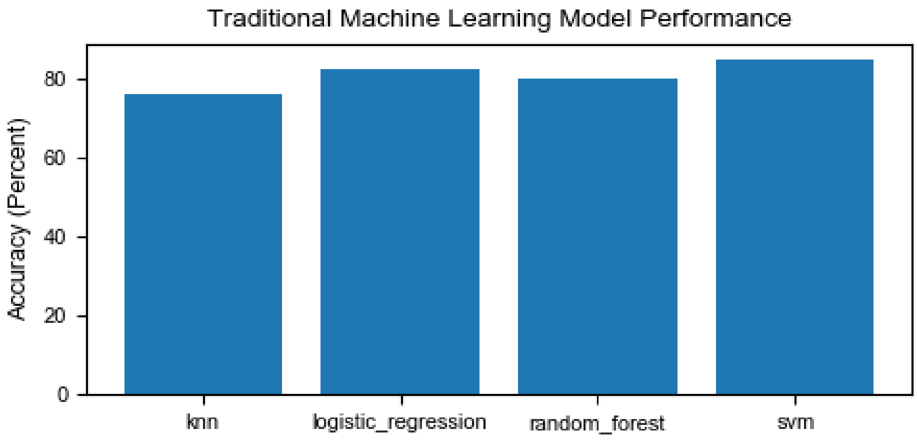


**Figure S3:** Validation set accuracy for each model

### Supplementary Note 6

#### Deep learning model creation

Sequential models were built using Tensorflow version 2.3.1. 1D CNN, LSTM, GRU, and CNN-GRU models were tested. Architecture specifics are given in GitHub. The 1D CNN, LSTM and

GRU models use a padded sequences of aspect ratio, perimeter, area and deformability as features. Hyperparameters are optimized using test data. ROC AUC and accuracy are reported using the validation data.

### Supplementary Note 7

### CNN model trained to detect morphology

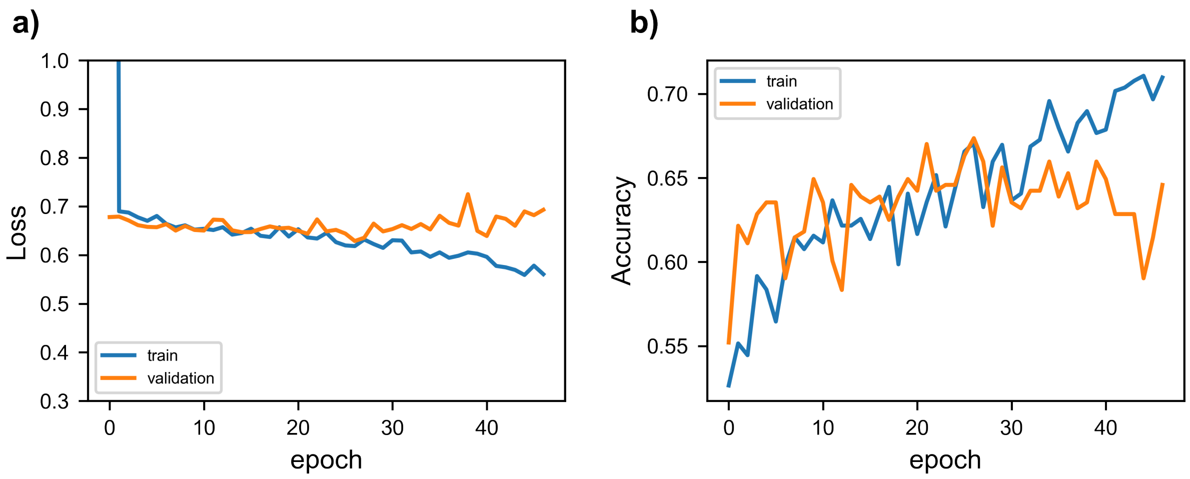


**Figure S4:** Validation set accuracy for each model.

### Supplementary Note 8

### CNN-GRU

Data were filtered, aligned and partitioned using the same methods described above for the GRU. Masks were cropped from the frame and padded to 90x90. A ellipse was fit to the cell and the two channels were concatenated. The optimizer used was RMSprop and the loss function was binary cross-entropy. The hyperparameters were optimized over 175 epochs using the test set. Once hyperparameter optimization was finished, validation accuracy was recorded and no further training was conducted. In order to further validate model performance, 5-fold cross validation was also conducted. Peak performances from each fold are listed below. The following model was used:

1. Conv2D - Filters 8

2. Conv2D - Filters 16

3. Conv2D - Filters 16

4. GRU - Length 50
5. GRU - Length 50
6. Dense - Length 1 - Activation: sigmoid

5-Fold Cross-Validation Results

| Fold 1 | Fold 2 | Fold 3 | Fold 4 | Fold 5 | Mean |
|---|---|---|---|---|---|
| 0.940 | 0.940 | 0.927 | 0.943 | 0.940 | $0.938 \pm 0.007$ |

#### Supplementary Note 9
#### 1D Convolutional Neural Network (CNN)

A 1D CNN was created and trained using Tensorflow 2.3.1. The optimizer used was RMSprop and the loss function was binary cross-entropy. The hyperparameters were optimized using the test set. Once hyperparameter optimization was finished, validation accuracy was recorded and no further training was conducted.

1. Conv1D - 16 Filters - Kernel Size of 10 - Activation: relu
2. Conv1D - 16 Filters - Kernel Size of 10 - Activation: relu
3. Conv1D - 16 Filters - Kernel Size of 10 - Activation: relu
4. Dense - Length 16 - Activation: relu
5. Dense - Length 16 - Activation: relu
6. Dense - Length 1 - Activation: sigmoid

#### Supplementary Note 10
#### Long Short-Term Memory Network

A LSTM network was trained using Tensorflow version 2.3.1. A model with the layers shown below was generated. The optimizer used was RMSprop and the loss function was binary cross-entropy. The hyperparameters were optimized using the test set. Once hyperparameter optimization was finished, validation accuracy was recorded and no further training was conducted.

1. Masking
2. LSTM - Length 50
3. LSTM - Length 50
4. Dense - Length 24 - Activation: relu
5. Dense - Length 1 - Activation: sigmoid

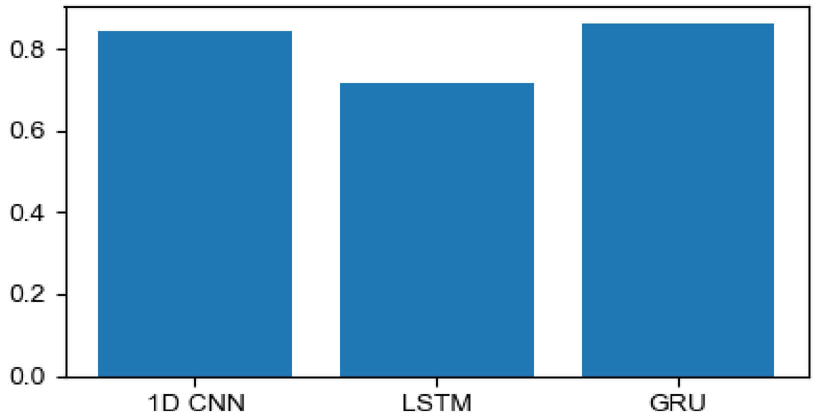


**Figure S5:** Accuracy is shown for each deep learning model trained using time-series features.

### Supplementary Note 11
### Gated Recurrent Unit Network

A GRU network was trained using Tensorflow version 2.3.1. A model with the layers shown below was generated. The optimizer used was RMSprop and the loss function was binary cross-entropy. The hyperparameters were optimized using the test set. Once hyperparameter optimization was finished, test accuracy was recorded and no further training was conducted.

1. Masking
2. GRU - Length 50
3. GRU - Length 50
4. Dense - Length 24 - Activation: relu
5. Dense - Length 1 - Activation: sigmoid

### Supplementary Note 12
### Detection, Segmentation and Tracking

A 4-layer CNN was created with each Conv2D layer containing a relu activation function, 3x3 kernel and 16 filters. The final layer was a single dense node with the sigmoid activation function. RMSprop was used to optimize the binary cross-entropy loss with a base learning rate of 0.0001. Roughly 9000 total images were used as inputs with 4000 background images and 5000 cell images. The input images were resized with padding down to 256x256. Pixel values were scaled down from 16 bits to 8 bits using the formula below.

$$\text{new frame} = 255 \times (\text{frame - min(frame)})/(\text{range(frame)})$$

After frames have been labeled by the CNN prefilter they are passed into the trained Mask RCNN where fits are obtained. Fits with a pixel area less than 600 or greater than 2500 are thrown out. The algorithm will then take data from the cell such as x-position, frame time and the binary mask to a dictionary with active events. The new mask data is appended to a list of a given event's mask data based on the new mask's change in frame time and x-position. If there are no cells that match the incoming data, the incoming data is assigned a new key in the dictionary. After the algorithm attempts to match incoming data with existing events, it checks to see if all incoming data has been assigned. If it has not, all incoming masks that have not been assigned are matched to the closest event. At the end of the video, active events are closed off. Events are only saved if there are more than 20 frames in the list.
